# CryptoBank: A Resource for the Identification and Prediction of Cryptic Sites in Proteins

**DOI:** 10.1101/2025.04.23.650184

**Authors:** Pedro Febrer Martinez, Thorben Fröhlking, Alberto Borsatto, Francesco Luigi Gervasio

**Author notes:** These authors contributed equally as first authors.

## Abstract

Cryptic binding sites in proteins, which are hidden in the absence of a ligand, offer opportunities to modulate targets previously considered ‘undruggable’. However, the scarcity of experimentally validated examples limits the development of predictive tools. Here, we introduce CryptoBank, a large-scale database of cryptic sites identified by applying a machine learning model to detect ligand-induced conformational changes in over 5.5 million structural alignments of unbound (apo) and bound (holo) protein pairs from the Protein Data Bank (PDB). Our analysis reveals that cryptic pockets are widespread, occurring in approximately 16.3% of protein clusters. Leveraging this resource, we fine-tuned a protein language model (PLM) to predict cryptic sites directly from protein sequence information. This sequence-based model achieves high precision (PR AUC 0.8) when query sequences share more than 20% identity with CryptoBank entries. Critically, we demonstrate its broader utility by predicting a cryptic site in human TPP1, a protein with less than 20% sequence identity to any CryptoBank entry, and validating its opening using molecular dynamics simulations. CryptoBank and the predictive PLM are publicly accessible via a web server, providing valuable resources for cryptic site discovery and drug development.

## 1 Introduction

Historically, pharmaceutical research has targeted clearly defined protein pockets, such as enzyme active sites, ion channels, and, more recently, well-characterized allosteric sites.^1–3^ In contrast, proteins lacking an obvious binding site in their native structure or binding with high affinity to their physiological substrate have often been classified as undruggable by conventional drug discovery methods.

Recent studies, however, have challenged this paradigm by demonstrating that many proteins, despite appearing to lack evident binding sites in their ligand-free (apo) states, can indeed bind small molecules through hidden ‘cryptic’ pockets.^4^ Cryptic pockets have therefore emerged as a valuable alternative for targeting challenging proteins, offering new binding sites that enable both direct inhibition and allosteric modulation.^5,6^

Despite their promise, the systematic identification of cryptic sites on proteins poses significant challenges. These pockets do not appear in experimentally determined unliganded structures but represent higher-in-energy states within the protein’s conformational ensemble^7–10^. The lack of readily available structural information complicates rational design and experimental identification, resulting in most cryptic pockets being serendipitously identified through high-throughput screening campaigns.

Computational methods offer a valuable alternative to cryptic sites identification. For instance, molecular dynamics (MD) based strategies and enhanced sampling algorithms have been widely used to sample the opening of cryptic cavities.^7,11–13^

More recently, machine-learning (ML) and mixed MD-ML approaches have also been developed for predicting the location of cryptic sites, with promising results.^14–16^ Additionally, the systematic application of some of these models to the human proteome, predicts that a substantial fraction of human proteins may harbour cryptic binding sites.

Despite their predicted abundance, the number of comprehensive, experimentally validated protein structure databases containing cryptic sites remains limited.^17^ Moreover, the only dataset that assess crypticity over an ensemble of binding site conformations includes no more than approximately 1500 structures.^18^ An ensemble-centric measure of crypticity is crucial, as it distinguishes pockets that rarely (if ever) form in the absence of a ligand from transient pockets that undergo rapid fluctuations and are less likely to serve as viable druggable sites.^19^

To address this challenge, we present here an integrated strategy to reveal cryptic binding sites in the PDB, CryptoBank, a structural database of cryptic pockets and a fine-tuned protein language model to reveal unknown cryptic pockets in protein targets. First, we develop a supervised machine learning strategy to quantify crypticity according to structural rearrangements between apo-holo configurations of a given binding site. We apply our scoring function to over 5.5 million structural alignments and find approximately 200 000 apo-holo-ligand combinations associated with crypticity, distributed across 1399 distinct clusters of 95% sequence identity. We then investigate whether sequence-level information of the identified cryptic sites can be extrapolated by a protein language model to predict novel cryptic pockets from apo protein sequences.

Protein Language Models (PLMs) have recently emerged as a powerful approach for protein structure and function prediction, and can be further fine-tuned for specific tasks.^20^ While fine-tuning has become standard practice in natural language processing, its application in protein modelling remains relatively novel, with promising results showing that task-specific and parameter-efficient fine-tuning can significantly enhance prediction precision across tasks involving protein structure, interaction, binding, stability and solubility. PLMs operate solely on sequence data. The major advantage is that protein sequences can be directly encoded and integrated into these models without complex preprocessing or extensive feature engineering.

We fine-tune the PLM for binary classification of binding site prediction at residue-level resolution. The model achieves state-of-the-art performance in predicting the location of cryptic binding sites in proteins and maintains high accuracy (ROC AUC) and precision (PR AUC) when applied to protein sequences that share at least 20% identity. However, the primary challenge remains in generalizing to entirely novel sequences not represented in the training data, highlighting the critical need for advancements in extrapolation techniques and robust detection of model hallucinations. Ultimately, we demonstrate with a relevant biomedical target, the telomere shelterin protein TPP1^21^, how the PLM predictions can be combined with state-of-the-art MD techniques to reveal novel cryptic binding sites, despite the TPP1 sequence sharing less than 20% identity with any CryptoBank entry.

## 2 Results

### Searching the PDB for cryptic sites

Cryptic pockets are typically defined as protein pockets that remain undetectable in the apo (ground) state of a protein but become readily identifiable in the holo state, where ligand binding reveals the cavity. Consequently, a valuable strategy to assess the crypticity of a given binding site is to compare its apo and holo states and quantify their structural differences. Building on these considerations, we designed a bioinformatics pipeline to identify apo-holo protein pairs from the PDB and assign a crypticity score to the detected binding sites. We started by considering all deposited protein crystal and cryo-EM structures with a resolution lower than 2.5 Å in the PDB (Fig. 1a). These structures were further filtered to retain only those with an associated UniProt accession and assigned to a 95% sequence identity cluster. The structures that met the filtering criteria were then split into individual protein chains and assigned to the apo or holo set. Initially, protein chains with a `nonpolymer_entity_count` of zero were classified as apo, while those with a `nonpolymer_entity_count` ≥ 1 were assigned to the holo set. As solvents used in structural assays may reveal cryptic sites, both sets were further refined based on a curated list of ions, low-mass compounds, and solvents (see *Methods*).^22^

**Figure 1:**
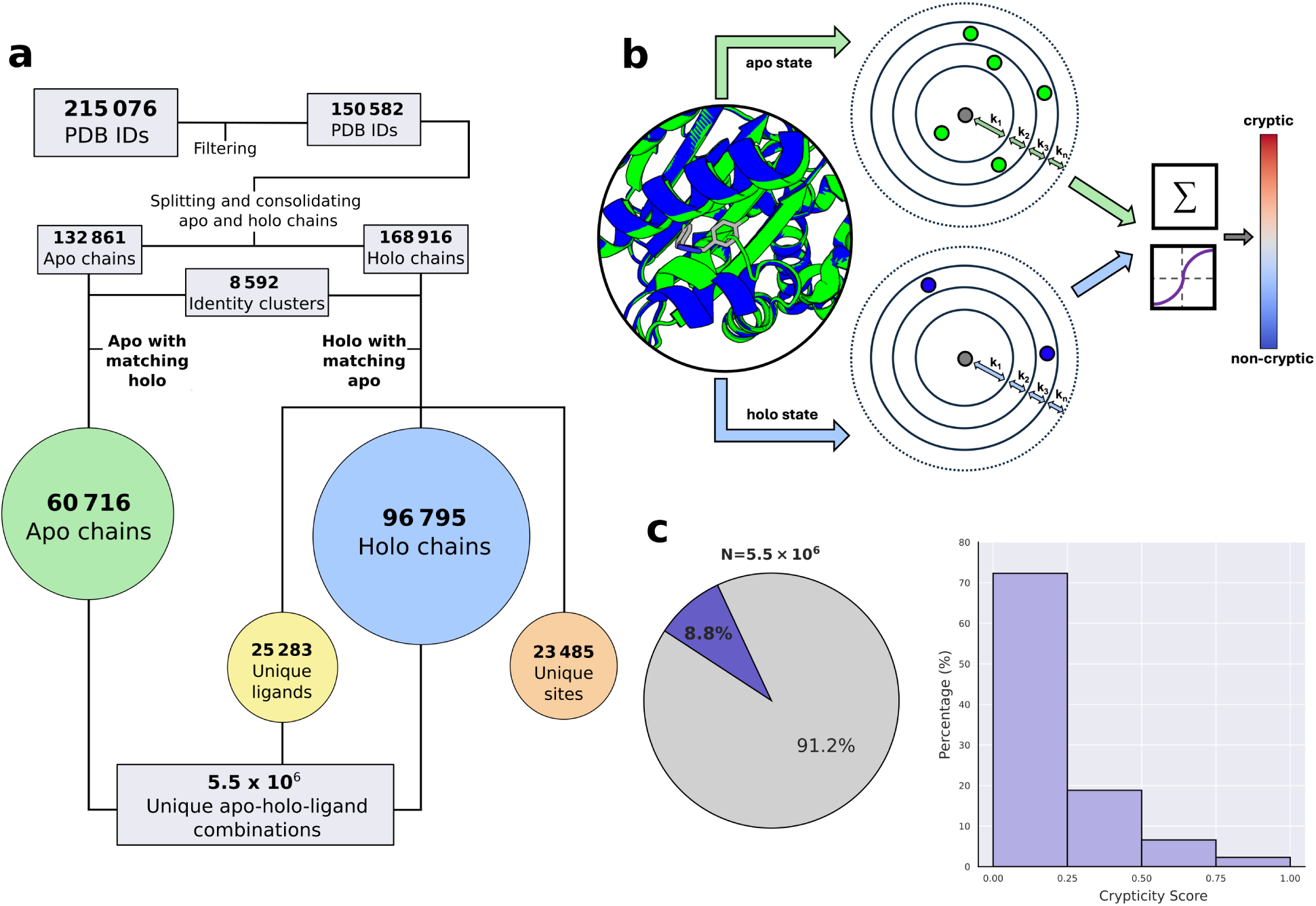
Computational pipeline and score distribution across the PDB. ***a)*** Bioinformatics pipeline to fetch and filter apo-holo-ligand combinations from the PDB. Starting from all X-ray and cryo-EM protein PDB IDs that match our filtering criteria, individual PDB IDs are then split into single protein chains and assigned to either the holo or apo set. Apo and holo chains sharing at least 95% sequence identity are clustered together and structurally aligned. Unique apo-holo-ligand combinations are stored in a data frame for subsequent scoring. ***b)*** Schematic of the scoring function. Starting from a structurally aligned apo-holo-ligand combination, the scoring function calculates the number of protein atoms within n concentric shells around each ligand atom for the apo and holo states separately. This is merged into a crypticity potential for the given apo-holo-ligand combination and normalized into a crypticity score (range 0–1) according to the number of atoms within each shell in the apo and holo states and the fitting parameters k. A score greater than or equal to 0.5 indicates crypticity. ***c)*** Pie chart of the crypticity scores obtained for all 5.5 million apo-holo-ligand combinations in CryptoBank (left), and histogram showing the distribution of these scores across the dataset (right).

We then matched apo and holo chains based on their sequence identity, grouping together chains that belong to the same 95% identity cluster in the PDB. Only protein chains with at least one apo and one holo chain within the same identity clusters were retained. This resulted in approximately 61000 and 97000 individual apo and holo chains, respectively, divided into 8592 95% identity clusters. Next, we identified all ligand binding pockets within each identity cluster. We selected a representative holo chain for each cluster and superimposed the remaining holo chains to this reference. We used a complete-linkage clustering algorithm to cluster together ligands whose centres of mass were within a 7 Å radius, resulting in 23485 unique ligand binding sites. Additionally, we assigned unique Tanimoto-based ligand identifiers to all ligands found in the different holo chains, ensuring the same chemical entities shared the same unique ligand identifiers, resulting in 25283 unique ligands. Following this filtering procedure, we structurally aligned each holo chain in a given cluster to all the apo chains in the same identity cluster. Additionally, each unique ligand was treated individually, so that each apo-holo pair was aligned once for every ligand present in the selected holo chain. For instance, if a cluster contained *n* apo chains, *m* holo chains, and *l* unique ligands, this resulted in a total of *n* × *m* × *l* intra-cluster alignments. Finally, only alignments that resulted in a *C_α_* RMSD lower than 2.5 Å were selected for scoring. Overall, this produced 5.5 million unique apo-holo-ligand combinations. Finally, we developed a supervised machine-learning model to assess the crypticity of the individual binding site conformations across the alignment set. As previously mentioned, cryptic sites undergo structural rearrangements to accommodate the ligand in the binding cavity. Specifically, the structural alignment of the apo and holo states of a cryptic site should result in atomic clashes between the ligand atoms and the apo residues, the severity and extent of which can be quantified via a scoring function (Fig. 1b). For a given apo-holo-ligand combination, the scoring algorithm determines the number of clashes between the ligand atoms and the aligned apo and holo states. If the orientation of the pocket-lining residues remains largely unchanged between the apo and holo structures, resulting in no atomic clashes, the site conformation is classified as non-cryptic; in contrast, significant clashes indicate a cryptic conformation of the site. Additionally, the model divides the ligand into segments to achieve higher spatial resolution of the binding region. For each segment, the model counts the number of protein atoms within *n* spherical shells of increasing radius centered on each ligand atom. Using this data, it calculates the probability that the segment is located in a cryptic region of the binding cavity. The final crypticity score for an apo-holo-ligand combination is defined as the average crypticity score across all ligand segments. The crypticity score is normalized between 0 and 1, with values of 0.5 or higher indicating crypticity. The training of the model involved tuning three hyperparameters, namely the number of ligand segments, the number of shells, and the L1 regularization term for the model weights (Fig. S1). A curated dataset of 199 apo-holo pairs, comprising 71 cryptic and 128 non-cryptic examples, was used for training. This dataset includes examples from prior studies as well as preliminary testing of our pipeline on the PDB (see Data availability).^14,15^ The model’s performance was evaluated using 5-fold cross-validation and validated on an independent test set, achieving a final accuracy of 89% (Fig. S1). We then applied the trained model to score the 5.5 million unique apo-holo-ligand combinations in our dataset (Fig. 1c). The resulting score distribution indicates that 8.8% of the alignments have a score of 0.5 or higher, suggesting potential site crypticity. To our knowledge, the resulting collection of unique apo-holo-ligand combinations exhibiting crypticity signals is the largest available to date.

### Quantifying crypticity of protein binding sites

To move from individual site conformation scores to an ensemble-level assessment of crypticity, we aggregated the crypticity scores of all alignments that map to the same binding site. The resulting site crypticity score represents the average crypticity score across all apo-holo-ligand combinations associated with the site.

Importantly, cryptic pockets are typically closed in the stable, unbound (apo) protein state. They form transiently within less stable, higher-energy conformations, which can be captured and stabilized by ligand binding. Since an apo protein structure represents a single conformation favoured by specific experimental conditions, variations in these conditions can yield different apo structures. This variability in the observed apo states can, in turn, affect the subsequent identification of cryptic sites.^18^ By averaging cryptic scores across multiple holo conformations of the binding site and different apo states, our approach accounts for these fluctuations, ensuring that a site is classified as cryptic only when a substantial fraction of its apo-holo comparisons are associated with higher crypticity scores. Following this scoring scheme, a binding site is flagged as cryptic if its mean site score is greater than or equal to 0.5.

Additionally, we considered the impact of different holo conformations on the site crypticity score. Chemically diverse ligands can map to different regions of a binding site, potentially influencing its crypticity. In some cases, multiple smaller ligands may bind to non-cryptic regions, while fewer, larger ligands expose cryptic regions of the site. To account for these scenarios, we implemented an outlier-oriented strategy to recover binding sites with a mean site score below 0.5 that still exhibit significant levels of crypticity (Fig. S2).

Given these considerations, 8.5% of the 23 485 binding sites in our dataset are classified as cryptic (Fig. 2a). In total, these 1985 cryptic sites correspond to 223 551 apoholo-ligand combinations with a cryptic score greater than or equal to 0.5, representing a two-order-of-magnitude increase over previous collections of cryptic site examples. Furthermore, these cryptic sites are distributed across 1399 distinct 95% identity clusters, accounting for 16.3% of all identity clusters currently in CryptoBank (Fig. 2b).

**Figure 2:**
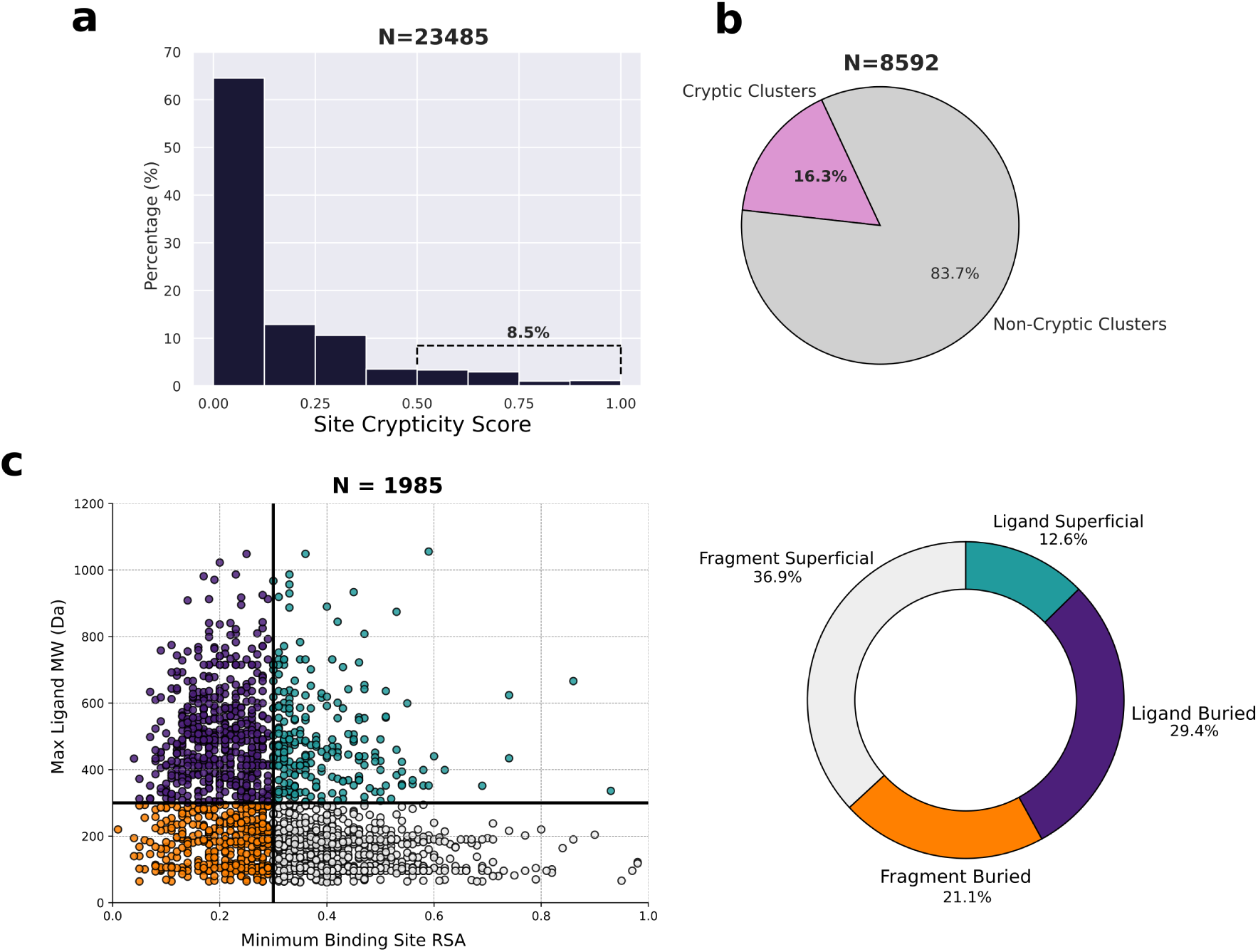
Site crypticity score and binding sites properties. ***a)*** Histogram of the site crypticity score distribution across all identified binding sites in CryptoBank. The score for each site is computed as the average of the individual crypticity scores across its conformations. For sites classified as cryptic by the outlier identification strategy, the mean score of the outlier conformations is reported. ***b)*** Pie chart showing the fraction of 95% sequence identity clusters that have a least one cryptic site in CryptoBank. ***c)*** Scatter plot of the minimum buriness and maximum ligand molecular weight for each cryptic site (left). Buriness is defined as the minimum relative surface accessible area (RSA) across the site’s conformations, while ligandability is defined as the maximum molecular weight of compounds binding to the site. Black lines indicate the thresholds used to classify sites as buried (RSA ≤ 0.3) and ligandable (MW ≥ 300 Da). Pie chart showing the fraction of cryptic sites classified as buried or superficial, and as binding either ligands (ligandable) or fragments (right).

Next, we calculated the molecular weight (MW) of the ligands binding to the identified cryptic sites, as well as the relative solvent-accessible surface area (RSA) of each site (Fig. 2c). Using these descriptors, we classified each site as either buried (RSA < 0.3) or superficial (RSA > 0.3) and determined whether it binds fragment-like (MW < 300 Da) or ligand-like (MW ≥ 300 Da) molecules. As a significant fraction of these sites bind more than a single ligand (Fig. S3), we selected the maximum ligand weight and minimum RSA associated with each binding region for our classification.

Our analysis indicates that 58% of cryptic sites are revealed by fragments, including both chemical fragments typically used in fragment-based screenings and small organic molecules commonly employed in crystallography techniques. This aligns with studies showing that small organic molecules, often present in buffers used during NMR and X-ray crystallography experiments, tend to accumulate at protein active sites and effectively map allosteric and cryptic sites in proteins.^23,24^ Of the cryptic sites exposed by fragments, 21.1% are located in buried regions of the protein, while the remaining 36.9% map to superficial areas of the protein surface. The presence of fragments in deeply buried regions suggests that some degree of protein rearrangement is required for these compounds to access the binding pocket, making these regions particularly promising for subsequent chemical design. Of the surface fragments, 12.8% and 5.6% bind to a surface patch targeted by at least one or two chemically distinct fragments, respectively. These regions may be close to a cryptic site, as studies have shown that protein regions interacting with multiple chemical probes are associated with crypticity.^9^

The remaining 42% of cryptic pockets can bind ligand-like molecules, most of which have molecular weights between 300 and 1200 Da. Specifically, 77% of these ligands fall within the rule-of-five-compliant range (300–500 Da). Interestingly, 70% of ligandbinding cryptic sites are located in deeper regions of the protein, while the rest are more superficial. Overall, among cryptic sites that bind at least one ligand in CryptoBank, 29.4% form buried pockets, whereas 12.6% are solvent exposed.

Finally, 16.4% of these ligandable sites also accommodate molecular fragments. Some of these examples highlight how fragment screening can initially identify cryptic pockets, which can later be exploited for ligand growth and optimization.

### Fragment library for cryptic pocket detection

Standard experimental and computational methods used for allosteric and cryptic pockets detection typically use small organic fragments to expose hidden cavities. The choice of organic probes to use is one of the key design challenges in such methods, with different fragments having varying effectiveness in mapping cryptic sites. Despite the huge efforts put into optimising these fragment libraries, most of the results obtained arise from either serendipitous findings or precisely designed libraries for one or at best few well characterised protein targets. In the context of cryptic pockets, additional challenges in designing such libraries can be attributed to the lack of a comprehensive collection of holo structures of cryptic sites. To address these challenges, we can leverage the data in CryptoBank to inform the design of tailored fragment libraries for cryptic pockets detection.

We started by dividing each ligand in our database into three distinct classes: ligands that bind exclusively to cryptic sites, ligands that bind solely to non-cryptic sites, and ligands that bind to both sets (Fig. 3a). Of the 25,283 unique ligands, 24% are found exclusively in cryptic sites, of which 63.8% have a molecular weight greater than 300 Da. The majority of ligands bind to non-cryptic sites (68.8%), while a minority fraction (7.4%) of ligands are found in both cryptic and non-cryptic sites. As expected, ligands targeting only one type of site tend to have a higher molecular weight, whereas those found in both types are typically fragment-sized. Heavier ligands are typically associated with higher levels of specificity for a binding region, suggesting that the majority of ligands targeting either cryptic or non-cryptic sites may exhibit some degree of selectivity.

**Figure 3:**
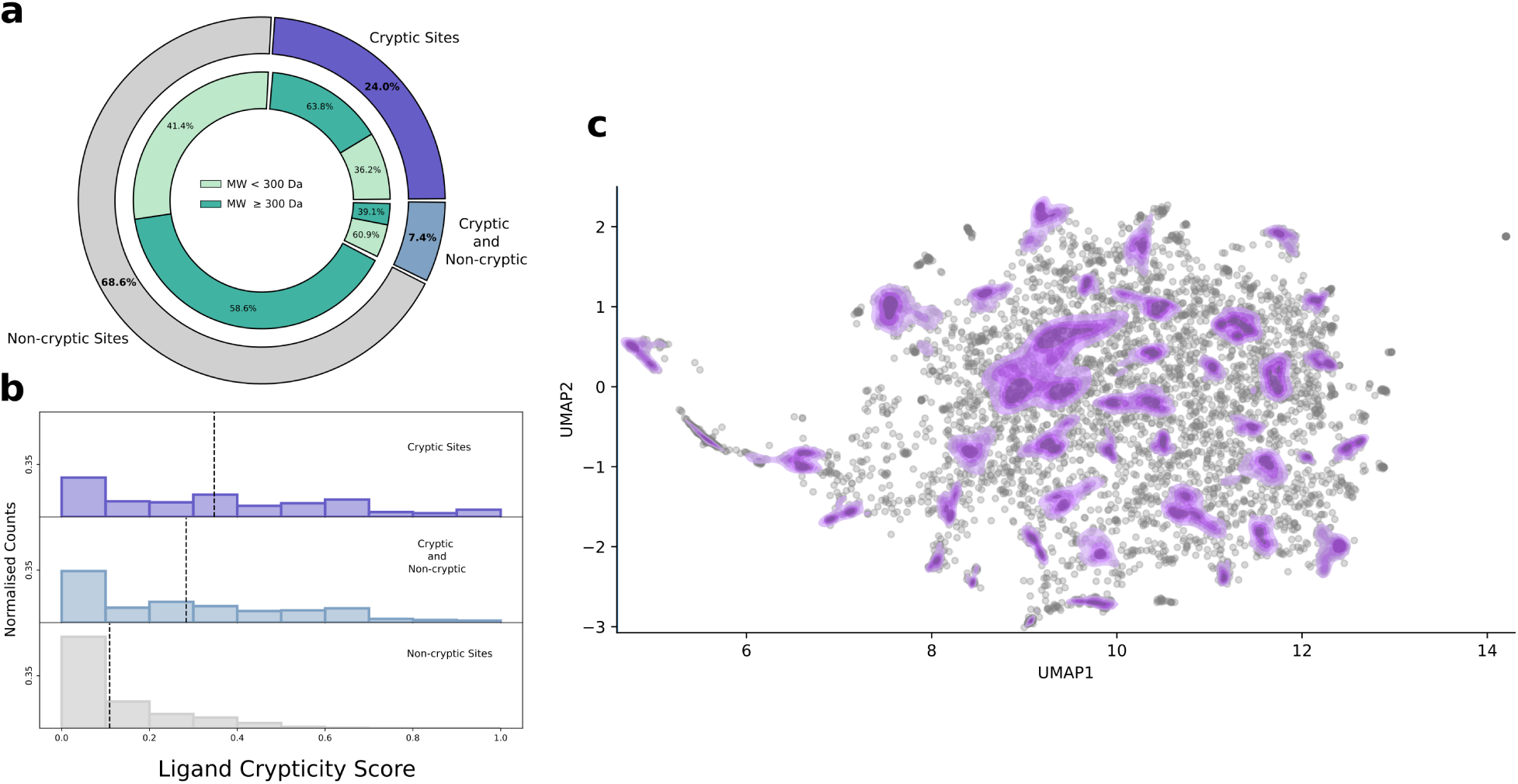
Ligand crypticity scores and the chemical space of compounds binding cryptic sites. ***a)*** Pie chart showing the distribution of ligands and fragments in CryptoBank binding to cryptic sites, non-cryptic sites, or both. The inner segments display, for each category, the proportion of compounds classified as ligand-like or fragment-like based on their molecular weight. ***b)*** Histograms of the ligand crypticity scores across all compounds in CryptoBank. From top to bottom: compounds found only in cryptic sites, in both cryptic and non-cryptic sites, and only in non-cryptic sites, respectively. Dashed black lines indicate the mean crypticity score within each class. Here, "ligands" refers to all compounds in CryptoBank, regardless of their molecular weight. ***c)*** Chemical space mapped by compounds found exclusively in cryptic binding sites. Each compound corresponds to a point in the UMAP projection. Clusters of points corresponding to high-density regions are grouped using HDBSCAN and highlighted in purple, while grey points represent lower-density regions that are not assigned to any cluster. Density lines for each cluster are obtained using Kernel Density Estimation (KDE). Only compounds with a molecular weight of maximum 1200 Da are considered for the analysis.

We then assigned each ligand a crypticity score (Fig. 3b). The ligand crypticity score is computed as the mean score of all apo-holo-ligand combinations for a given ligand. Plotting the distribution of the obtained ligand crypticity scores confirms that, on average, ligands that bind cryptic sites induce greater structural rearrangements of the binding region (higher score). In contrast, ligands exclusively targeting non-cryptic sites are associated with lower ligand crypticity scores, while ligands present in both exhibit an average score falling between the two classes.

Finally, we clustered the 6000 ligands based on Morgan fingerprint similarities (Fig. 3c). We identified similarity clusters in the UMAP space using HDBSCAN to detect local densities, resulting in 38 distinct clusters. The molecule closest to the centroid of each cluster was selected as the cluster representative (Fig. S4). When computing the Tanimoto similarities between these cluster representatives, we obtained a maximum similarity of 0.2, indicating strong chemical diversity within the set (Fig. S5). This library can serve as a starting point for refining fragment libraries used to screen for cryptic pockets.

Additionally, the data in CryptoBank can be leveraged to generate fragment libraries tailored for specific proteins. Under the assumption that similar protein tend to bind similar ligands, one can, starting from the sequence of a target protein, extract ligands found in cryptic pockets of similar proteins.^25^ Moreover, the ligand crypticity score can be used as a ranking criterion to prioritize molecules for screening experiments. In cases where no cryptic sites are present, the general ligand list can still provide added value to inform screening campaigns for cryptic site detection.

### Cryptic sites in pharmaceutically relevant protein targets

To evaluate the relevance of cryptic pockets in high-priority protein targets, we crossreferenced the data in CryptoBank with the Open Targets platform.^26^ The Open Targets platform leverages publicly available datasets to systematically evaluate and prioritize drug targets. By integrating large-scale experimental data with existing biological and clinical knowledge, the platform constructs and scores associations between potential targets and diseases.

We selected all protein-coding genes in the Open Targets dataset with an association score exceeding 0.5 for at least one human disease, resulting in a total of approximately 6000 disease-associated genes. We then collected the UniProt accession numbers associated with the selected gene set and mapped the Uniprot accession numbers to the sequence identity clusters in our database. Following this procedure, we identified 866 distinct 95% identity clusters in CryptoBank associated with human disease. Among these 866 proteins, 31.1% harbour a cryptic site (Fig. 4a). These findings are in line with previous estimates on the abundance of cryptic sites in proteins of pharmaceutical interest.^14,15^

**Figure 4:**
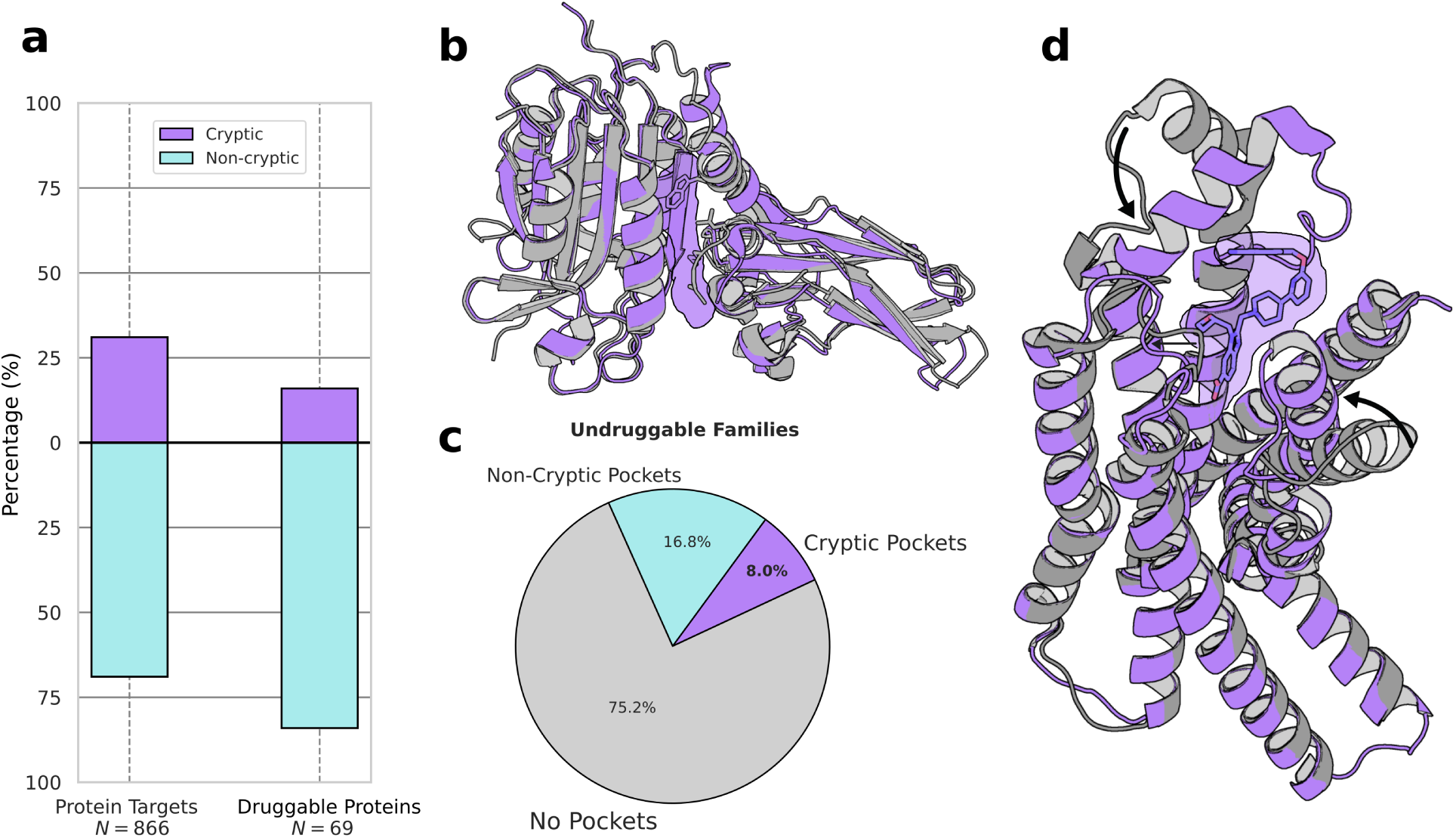
Crypticity of protein targets in the Open Targets platform. ***a)*** Bar plot showing the percentage of protein targets and druggable proteins harbouring cryptic sites. The set of druggable targets includes proteins with only one binding site. ***b)*** Structural alignment of MALT-1 in its apo (gray, PDB: 3V55) and holo (purple, PDB: 7AK0) states.^27,32^ The cryptic cavity observed in the holo structure is shown as a surface, the ligand is omitted for clarity. Protein residues that occlude the cryptic pocket in the apo structure are represented as sticks. ***c)*** Pie chart of proteins in CryptoBank belonging to undruggable families and harbouring at least one ligandable pocket. The fraction of proteins with at least one ligandable cryptic pocket is shown in purple, while proteins with ligandable non-cryptic pockets and those with no ligandable pockets are shown in cyan and grey, respectively. ***d)*** Structural alignment of GLP1R in its apo (gray, PDB: 6X18) and holo (purple, PDB: 6X1A) states.^33^ The cryptic cavity observed in the holo structure is shown as a surface, and the ligand as sticks. Conformational rearrangements required to form the cryptic cavity are highlighted with arrows.

Additionally, we investigated how many of the disease-associated proteins in CryptoBank are targeted by at least one approved drug or are involved in advanced clinical trials. We focused exclusively on therapeutic strategies involving small-molecule binders and PROTAC-like compounds, as both modalities require a binding pocket in the protein target. Furthermore, to increase the likelihood that the binding site targeted by these compounds corresponds to the binding site in our database, we restricted our search to proteins in CryptoBank with only one ligand-binding site. Surprisingly, 16% of the 69 identified target pockets meeting these criteria are cryptic. For instance, the mucosa-associated lymphoid tissue lymphoma translocation protein 1 (MALT-1) clearly exposes a cryptic allosteric pocket forming in the caspase domain of the protein (Fig. 4b)^27^. This cryptic site is currently under investigation in a phase 1 clinical trial (NCT05544019) as a promising target for treating patients with relapsed or refractory B-cell neoplasms.

Next, we checked how many of the proteins in CryptoBank present a ligandable cryptic pocket and are assigned to undruggable families in the OpenTarget database. Typically, pharmaceutically relevant binding sites are located in buried regions of the protein and accommodate larger ligands, therefore we focused our analysis only on proteins with pockets with these characteristics. Specifically, we selected binding pockets with an RSA < 0.3 and that bind ligands with 25-50 heavy atoms. The results show that of the proteins considered undruggable in CryptoBank, 24.8% have a buried, ligandable site, with 8% of these proteins harbouring potentially druggable cryptic pockets. Although preliminary, these results indicate how cryptic pockets offer valid options for expanding the druggable proteome, as the presence of a targetable pocket is essential for small-molecule–mediated drug discovery.

Finally, we cross-referenced CryptoBank with a structural database of G proteincoupled receptors (GPCRs), namely GPCRdb.^28^ Currently, CryptoBank contains 34 GPCR structures distributed across five distinct 95% identity clusters. Due to the stringent filtering criteria applied when querying the PDB, only a small fraction of the available GPCR structures are included in the database. Nonetheless, we identified a cryptic binding site in the Glucagon-Like Peptide 1 Receptor (GLP1R) (Fig. 4c). The structural alignment of the receptor’s apo and holo states clearly shows the formation of a cryptic pocket around the ligand, resulting from the partial unfolding of an alpha-helix, the displacement of a loop, and the movement of a second alpha-helix toward the binding site. This observation aligns with mounting evidence that GPCRs exhibit varying levels of crypticity, with some cryptic sites being directly targeted in drug discovery campaigns.^29–31^

### PLM Fine-Tuning for cryptic binding sites

In Fig. 5 we collected the evaluation for the fine-tuned Prot-T5-XL-UniRef50 model performance across training, validation, and test set on exclusively cryptic systems.^34^ The ROC curve for the training set yields a high AUC of 0.98, because of the high data imbalance with a positive class ratio of 0.07 making it easier for the model to achieve high AUC. We select the best model as the one that is maximizing the cross-entropy loss on the validation set yielding an AUC of 0.96. Since the validation data share a similar imbalance with a positive class ratio of 0.06, the ROC AUC may overestimate the extrapolation behavior to systems not seen during training. Benchmarking against the test set results in an AUC of 0.74 for the ROC curve (Fig. 5a), indicating that unique sequences impose a significant challenge for the PLM. To highlight the model’s ability to achieve high precision, an important aspect of binding site prediction due to its highly imbalanced nature, with the positive class comprising at most 7% of all labels, we compute the model’s precision-recall curves (Fig. 5b). According to this metric, the various datasets differ even more significantly in terms of their AUCs. The precision on the training data is high with a PR AUC of 0.88 corresponding to an increase in precision of more than an order of magnitude compared to a random model. Comparing to the precision of the selected model with respect to the validation set a slight decrease in PR AUC to 0.80 can be seen. This shows that the PR AUC is the more adequate metric for imbalanced prediction tasks. Importantly the model retains significant precision indicating it is capable of extrapolation to new systems with still an order of magnitude higher precision than a random prediction. Benchmarking against the test set yielded an AUC 0.17 for the precision-recall curve. While the high accuracy obtained on the test set can be misleading for imbalanced data the PR AUC instead reveals the significant drop in performance. The retained precision corresponds to a 3-fold increase in precision compared to a random model. The computed PR AUC for the PLM reveal its limitations, but also demonstrates that a protein language model fine-tuned on CryptoBank can precisely predict cryptic sites from apo protein sequences, achieving a PR AUC of 0.8 when the query sequence shares more than 20% identity with a CryptoBank entry. Therefore the PLM can be considered among the state-of-the-art models for predictive models applied in the context of cryptic and non-cryptic protein-ligand interaction^14,15,35^, underscoring that signal about cryptic binding site locations is encoded at the sequence level. Furthermore, the model demonstrates a limited but notable ability to extrapolate signals to unique sequences, emphasizing its generalization capability beyond the training data.

**Figure 5:**
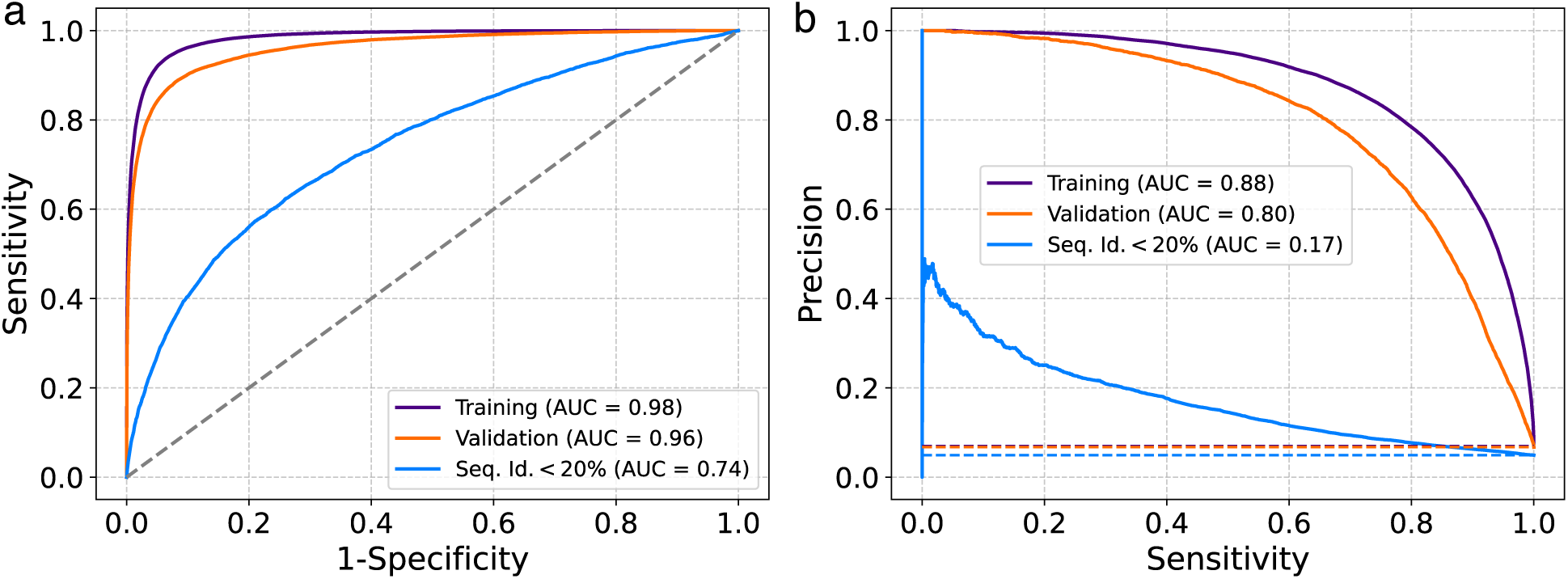
Performance of the ‘Prot-T5-XL-UniRef50’ model after fine-tuning evaluated on training, validation and test set for cryptic protein sequences only. *a)* ROC curves showing AUCs of 0.98, 0.96 and 0.74 respectively, while random prediction will lead to an AUC of 0.5. *b)* Precision-recall curves show AUCs of 0.88, 0.80 and 0.17 respectively, demonstrating high precision for the imposed imbalanced binary classification task. Performance of random predictions corresponds to the fraction of the positive class and is maximally at 0.07 for all sets visualized as dashed lines with matching colors.

### PLM prediction applied to TPP1

We tested an application of our fine-tuned PLM model to support screening efforts to identify novel cryptic pockets. To this end, we selected the telomere shelterin protein TPP1, as no small molecule-bound structures are currently available and therefore not present in the training sequences used to fine-tune the PLM model. TPP1 plays a critical role in the regulation of telomere length by both recruiting and activating telomerase, making it a highly interesting therapeutic target.^36^ Notably, the PLM predicts the presence of an allosteric binding site far from the protein-protein interface typically formed between TPP1 and telomerase (Figure 6a). Qualitative visual inspection of the PLM predictions reveals a cluster of residues with a high probability of forming a cryptic binding site, located on an *α*-helix and its connecting loop.

**Figure 6:**
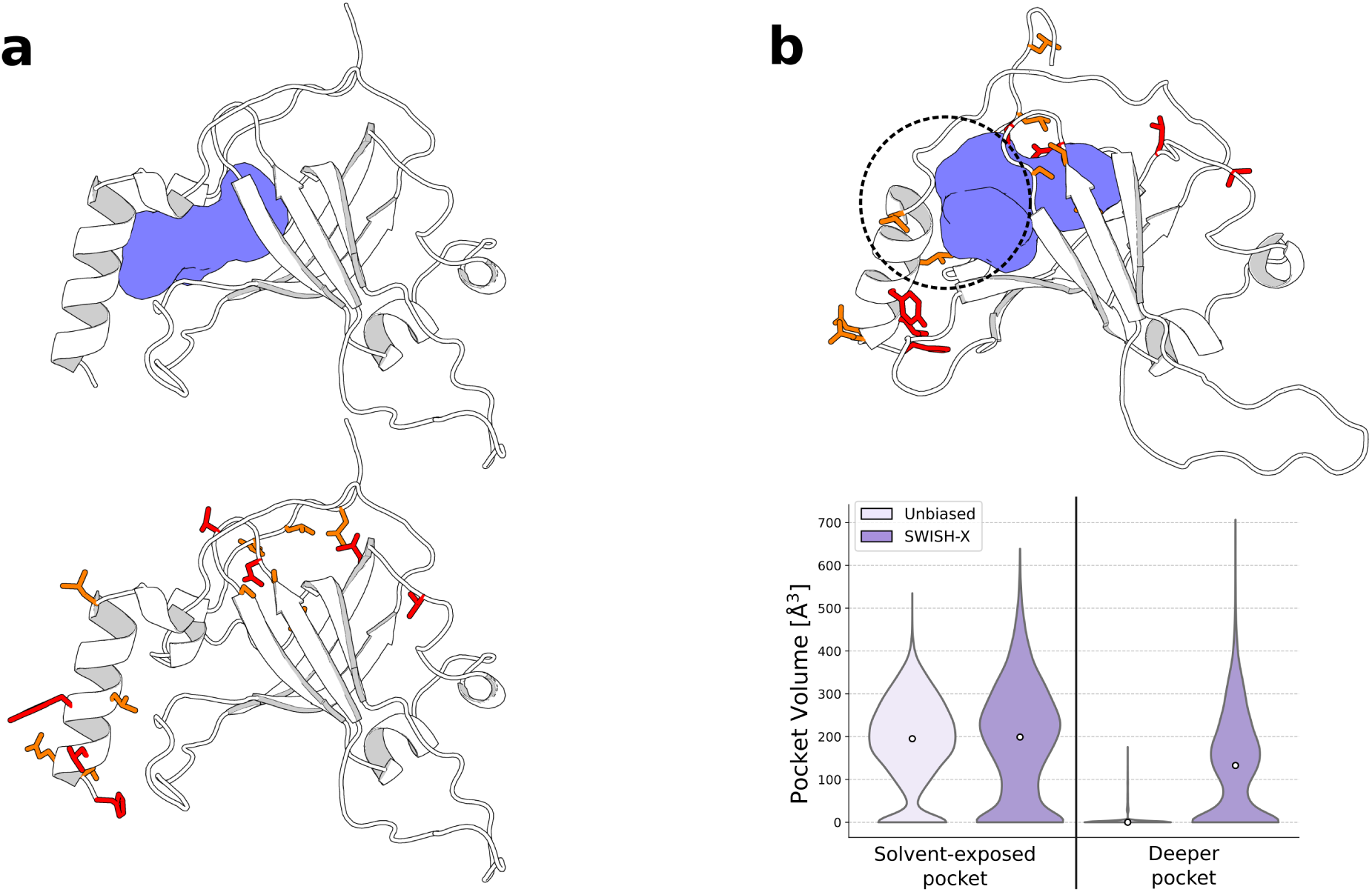
Predicted cryptic pocket in TPP1. ***a)*** Structural representation of TPP1 (top panel). The internal cavity identified in the cryo-EM structure is displayed as a purple surface (PDB: 7TRE). The bottom panel shows the crypticity prediction from the finetuned PLM for TPP1: residues with a normalized score between 0.4 and 1 are rendered as sticks, with residues scoring above 0.6 highlighted in red and those scoring between 0.4 and 0.6 in orange. ***b)*** Opening of the cryptic cavity connecting the surface of TPP1 with its core (top panel). The dashed circle indicates the major conformational changes required to fully expose the cavity, which is depicted as a purple surface. The bottom panel shows violin plots representing the volume of the identified cryptic pocket along the simulations.

Interestingly, the cryo-EM structure of the TPP1–telomerase complex reveals the presence of a buried internal cavity within the TPP1 core (PDB ID: 7TRE).^21^ However, it remains unclear how this cavity could become solvent-accessible. We therefore hypothesised that, as predicted by our fine-tuned PLM model, the opening of a transient cavity in this region could act as an entry point to deeper pockets within the protein core. To test this hypothesis, we first performed an unbiased MD simulation to monitor pocket formation. Notably, the predicted region formed a solvent-exposed cavity during the simulation, while the internal cavity quickly closed (Fig. 6b). We then ran SWISH-X simulations^12^ to investigate whether the transient cavity could connect the deeper cavity observed in the cryo-EM structure to the protein surface. Remarkably, we recovered fully connected conformations of both the surface-accessible and internal cavities, with the entry point in direct contact with the predicted cryptic region. Specifically, to fully expose the cavity, the *α*-helix containing most of the predicted cryptic residues tilts relative to its position in the crystal structure. Additionally, the loop connecting this helix to the central region of the protein, shifts outward, linking the deeper pocket to the protein surface.

Overall, these results demonstrate that fine-tuning with cryptic binding examples enables the PLM model to successfully predict cryptic regions that open during simulation, highlighting its potential to inform drug discovery efforts.

## 3 Discussion

The challenge of targeting proteins lacking well-defined binding pockets in their native state has significantly limited the scope of drug discovery. Cryptic sites, which form upon ligand binding or through transient protein fluctuations, represent a promising but underexplored avenue for addressing these "undruggable" targets. However, progress has been hampered by the scarcity of systematically identified and validated cryptic sites. This study introduces CryptoBank, a large-scale, computationally derived database of cryptic binding sites, and demonstrates the potential of leveraging this resource for predictive modeling. Our primary contribution is the creation of CryptoBank, populated by analyzing over 5.5 million structural alignments between apo and holo protein states from the PDB. Using a supervised machine learning model trained to quantify ligand-induced structural rearrangements, we identified approximately 200,000 apo-holo-ligand combinations exhibiting cryptic characteristics. These aggregate into 1,985 unique binding sites classified as cryptic, distributed across 1,400 distinct protein clusters (16.3% of clusters analyzed). This represents a significant expansion, by potentially two orders of magnitude, over previously available datasets of cryptic sites. The scale and the ensemble-based nature of our crypticity assessment, which averages scores over multiple apo and holo conformations, provide a more robust view compared to single-structure analyses, accounting for the inherent conformational flexibility influencing site formation. CryptoBank offers immediate value as a resource for drug discovery and biology. Firstly, it provides experimentally-grounded structural information on pockets that are often invisible in apo structures, offering starting points for structure-based design against difficult targets. Our characterization of these sites based on ligand molecular weight and solvent accessibility revealed diverse scenarios: 58% are revealed by fragment-like molecules, while 42% bind larger, ligand-like compounds. Notably, a significant portion (70%) of ligand-binding cryptic sites are buried, aligning with characteristics often sought for druggable pockets. The finding that cryptic sites exist in proteins previously deemed "undruggable" (with 8% of such proteins in our dataset showing a potentially druggable cryptic site) directly addresses a major bottleneck in expanding the druggable proteome. Furthermore, the identification of cryptic sites in therapeutically relevant targets like MALT-1 and GLP1R, and within 31.1% of disease-associated proteins from Open Targets present in our dataset, highlights the immediate applicability of CryptoBank to ongoing drug discovery efforts. The curated list and clustering of 6,000 ligands found exclusively in cryptic sites also provide a foundation for designing tailored fragment libraries optimized for cryptic site detection. Beyond serving as dataset, we demonstrated that the sequence-level information inherent in CryptoBank can be harnessed for prediction. By fine-tuning a protein language model on our dataset, we developed a tool capable of predicting cryptic site propensity directly from protein sequences. When the query sequence shares more than 20% identity with a CryptoBank entry, high prediction accuracy and precision of AUC 0.96 and PR AUC 0.80 can be achieved. The biggest challenge remains in generalizing to entirely novel sequences, as reflected in the performance drop to test set AUC 0.74 and PR AUC 0.17. Importantly the successful prediction and subsequent simulation-based validation of a potential cryptic site opening in TPP1, a protein not present in the training data and sharing less than 20% identity with any CryptoBank entry, serves as a compelling proof-of-concept. It shows how the PLM can guide resource-intensive computational or experimental methods (like MD simulations or fragment screening) towards regions most likely to yield novel binding pockets. However, certain limitations should be recognised. The definition and identification of cryptic sites is inherently dependent on the structural data available in the PDB, which may be biased towards certain protein states or families. While the performance of the PLM is promising, it suggests that prediction of cryptic sites in proteins that are highly dissimilar to the training set remains a challenge, requiring further advances in model architecture or training strategies, possibly integrating structural information. In conclusion, CryptoBank provides the largest curated collection of cryptic binding sites to date, offering a valuable resource for understanding the structural basis of cryptic site formation and for initiating drug discovery campaigns against challenging targets. The associated PLM, despite limitations in generalization, represents a significant step towards sequence-based prediction of these elusive pockets. Together, CryptoBank and the predictive model offer powerful tools to systematically explore cryptic sites, expanding the landscape of druggable targets and potentially unlocking new therapeutic strategies for previously intractable diseases.

## 4 Methods

### Data collection and filtering

We obtained protein structures from the RCSB Protein Data Bank (PDB) as of 2025-03-12, using a filtering approach via the RCSB Search API. The filtering criteria were implemented via the rcsb-api python package.^37^ Specifically, the following filtering criteria were used: (1) a resolution filter to select structures with resolution higher or equal to 2.5 Å, (2) a preliminary holo structure filter selecting entries with at least one nonpolymer entity, (3) an apo structure filter selecting entries with no non-polymer entities, and (4) a cluster membership filter ensuring all structures belong to a sequence identity cluster. These criteria were then combined using logical operators to create the final queries for holo and apo structures. Following initial structure identification, we used the RCSB Data API to extract information for each identified structure ID using a predefined JSON file containing the fields of interest. A custom Python function was used to parse the JSON responses and organize the data into a structured dataframe format. From each structure ID, individual protein chains were extracted separately, preserving chain-specific information including associated ligands. For holo structures, chains were further subdivided based on their bound ligands, creating individual entries for each unique chain-ligand combination. Protein chains from holo structure IDs that lacked associated ligands were reclassified as apo chains. For holo chains with ligands, we implemented an additional filtering step to exclude non-relevant ligands based on an exclusion list containing ions, low molecular weight compounds (< 60 Da), solvents, and other non-functional molecules. The molecular weight of each ligand was determined using the formula_weight parameter from the PDB API. Holo chains containing only excluded molecules were also reclassified as apo chains. During this filtering process, we also removed chains lacking UniProt identifiers (typically representing non-protein polymers such as DNA, RNA, or small peptides) and chains without a cluster_id_95 assignment (95% sequence identity clusters). To identify and retain only valid apo-holo pairs, we matched protein chains sharing the same cluster_id_95, indicating 95% sequence identity. For each holo chain-ligand combination, we identified all available apo counterparts within the same sequence cluster. This pairing approach resulted in approximately 5.5 million apo-holo pairs for subsequent structural alignment and cryptic pocket identification.

### Structural alignment

Prior to alignment, alternate atomic locations in both protein chains and ligands were removed to maintain a single set of atomic coordinates for each protein residue and ligand. For each apo-holo pair, we performed a global alignment of the apo structure to the holo structure using PyMOL’s alignment algorithm with 5 refinement cycles. Following this initial alignment, binding site residues were identified within a 4 Åradius around each ligand in both the holo and aligned apo structures. The complete set of resolved residues in each structure was also saved for subsequent analysis. The aligned coordinates of the holo chain, apo chain, and ligand were saved in XYZ format, and the RMSD between the aligned structures was computed to quantify structural differences. To improve alignment quality in the binding site region, we performed 10 additional local alignments with progressively increasing selection radii (ranging from 10 to 20 Å) around each ligand. The local alignment yielding the lowest RMSD value was selected as the optimal alignment for each apo-holo-ligand combination, and the corresponding coordinates were saved in XYZ format. The so obtained XYZ files are the input of our scoring function, corresponding to apo-ligand and holo-ligand distance matrices.

### Scoring function

Starting from a structurally aligned apo-holo pair, our supervised machine learning model takes apo-ligand and holo-ligand distance matrices as input and assigns a probability to the site conformation to be cryptic. The distance matrix, *d*, contains pairwise distances between ligand and protein atoms, calculated within a 4 Å cut-off.

To quantify potential clashes, the distance matrix is segmented into *N* shells, each defined by a distance range [*d_n_*, *d_n_*_+1_). For each shell *n*, a potential *V* is computed by summing the indicator function over the distances falling within the shell and weighting it by a fitting parameter *k_n_*:

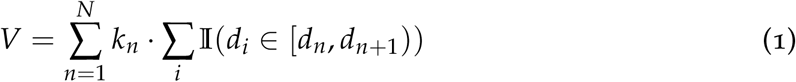

where *k_n_*is a learned parameter specific to shell *n*, *d_i_* represents the individual distances, and I(*d_i_*∈ [*d_n_*, *d_n_*_+1_)) is an indicator function that equals 1 if *d_i_* is within the shell’s range and 0 otherwise.

The potentials from the apo and holo states are aggregated to compute the total energy *E*_system_, which is averaged over the number of distances in these states as follows:

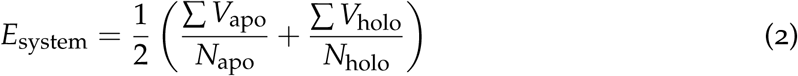

where *V*_apo_ and *V*_holo_ are the potentials from the apo and holo states, and *N*_apo_ and *N*_holo_ are the number of distances in these states. The computed energy *E*_system_ is then transformed into a probability *P* using a sigmoid function. This probability reflects the likelihood of superpositions or clashes, with higher probabilities corresponding to stronger superpositions.

In order to allow the model to systematically account for contributions from different regions of the ligand-protein interface the formalism above can be generalized by dividing the ligand and its surrounding protein atoms into *S* segments. For each segment *i*, the energy *E*_segment,*i*_ is computed independently. The probabilities for each segment are aggregated to produce a final probability *P*_segment_ as follows:

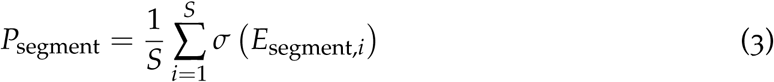

where *σ* is the sigmoid function, and *E*_segment,*i*_ is the energy of segment *i*.

### Training of the scoring function

We start by manually curating a new dataset as a combination of systems considered relevant in the context of cryptic and non-cryptic binding protein systems in previous works resulting in a collection of 199 systems of which 71 are labelled cryptic and 128 are non-cryptic.^14,15^ We train the scoring function, i.e. a binary classifier, on this new dataset of systems using the algorithm described previously and binary cross-entropy loss.

In order to avoid overfitting, we perform 5-fold cross-validation and apply L1 regularization to the loss function in the form 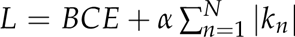 where *BCE* represents the binary cross-entropy loss, *k* are the free parameters of the classifier model, and *α* is the regularization strength controlling the penalty on the magnitude of the parameter values *k*. The absolute value |*k_n_*| applies L1 regularization, which encourages sparsity in the parameter values effectively prioritizing contributions from certain regions of the ligand-protein interface over others.

We optimize three hyperparameters by evaluating model performance across multiple dimensions. First, we iterate over classifiers using distance matrices derived from splitting the ligand into increasing numbers of segments, characterised by the number of segments *S*. For each segmentation value, we perform a grid search over the number of shells *N* and the regularization strength *α*. This combined approach allows us to identify the optimal number of shells *N* and regularization strength *α* for the respective segmentation *S*. We retrain the classifier 30 times, using optimal hyperparameters on the full dataset. For each iteration, we randomly split the data into training and validation sets, varying the validation set size, and minimize the binary cross-entropy loss. The best performing model was found to have the following parameters: S=3, N=4, and *α* = 1.6 × 10^−5^.

### Binding site identification

We developed a Python script to label binding sites within protein clusters, using ligand positions across all holo protein chains in a given cluster. Initially, all holo chains in the cluster are aligned to a reference holo chain using PyMOL. Subsequently, the center of geometry (COG) is computed for each ligand. A complete-linkage clustering algorithm then groups ligands with COGs within a 7 Å radius. This ensures that each ligand is assigned to a single cluster (binding site), with all ligand COGs within a cluster being within a 7 Å radius of each other. Finally, unique binding site labels are assigned to each identified cluster, representing distinct binding sites within the same cluster.

### RSA and ligand properties

We extract the relative per-residue solvent accessible surface area (RSA) using PyMOL’s get_sasa_relative function applied to the selection of residues within 5 Å from the ligand, the average RSA of the selected residues is used as the binding site RSA. Simultaneously, we determine the number of ligand atoms and the ligand’s molecular weight directly from the resolved structure. These structural properties are then compared against the complete ligand information provided by the PDB API, namely, the total atom counts and molecular weight. This comparison allows us to identify cases where the experimental structure might be incomplete or differ from the expected ligand structure.

### Datasets for PLM fine-tuning

We perform PLM fine-tuning for two different datasets. The dataset used for the first PLM fine-tuning in this study comprises exclusively cryptic holo protein structures. Sequences shorter than 50 amino acids were excluded to ensure high structural complexity, while sequences longer than 1500 amino acids were excluded due to GPU memory limitations when fine-tuning the PLM. We notice that the distribution of protein sequence lengths in the PDB is peaked between 100 and 300 amino acids. For each system, we retained the cryptic protein segments. Binding sites are defined as residues with at least one atom located within 5 Å of a cryptic ligand atom and translated into binary labels for the subsequent per-residue classification task. We compute sequence identity using MMseqs2^38^ and exclude a test set of 300 unique sequences that share less than 20% sequence identity with any other sequence. Subsequently, all remaining sequences are split 80/20 into training and validation sets.

The dataset used for the second PLM fine-tuning comprises holo protein structures available in the PDB. As in the first case, sequences are filtered by length (50-1500 amino acids), and binding sites are defined using a 5 Å distance threshold and converted to binary labels. However, for each system we retain all available protein segments rather than only cryptic regions. Additionally, we define valid ligands as those not containing nucleic acids, ions, or solvents, and not exhibiting more than 50 atoms. Starting from the PDB holo IDs provided by PLINDER^39^, we retain all sequences of the required length and bound to valid ligands. We compute sequence identity using MMseqs2 and exclude a test set of 645 unique sequences that share less than 20% sequence identity with any other sequence. The remaining sequences are scored using the probabilities of the binary classifier described in the previous subsection. Sequences with a score higher than 0.5 are assigned to the cryptic binding site-containing training set, and all remaining sequences with a sequence identity higher than 0.4 are also included in the training set. This results in over 48,000 sequences for training and over 90,000 sequences for validation. The results for this second fine-tuning experiment can be found in the SI.

### PLM fine-tuning

We refined ProtTrans’s PLM Prot-T5-XL-UniRef50 that was trained on 393 billion amino acids using a masked language modelling objective^34^. For hyperparameter choices we followed previous studies^20^. We set learning rate to 3 × 10^−4^ and choose batch size 8, achieved via batch size of 1 and gradient accumulation of 8. Mixed precision training was enabled to improve computational efficiency. We use Low-Rank Adaptation (LoRA), introducing trainable LoRA layers to the attention mechanisms. To increase classification performance, a convolutional neural network prediction head was integrated. This consisted of a 1D convolutional layer with 512 output channels and a kernel size of 3, followed by a linear layer mapping the output to the target number of classes. The model is trained to perform binary classification on residue level for 50 epochs, balancing computational efficiency and model convergence. We select the model with the lowest cross-entropy loss for the validation set. Models are trained and evaluated using an AMD Ryzen Threadripper PRO 5995WX CPU and a Nvidia RTX4090 GPU.

### MD and SWISH-X simulations of TPP1

The atomistic unbiased molecular dynamics simulations were performed using GRO-MACS 2023^40^ patched with PLUMED 2.9^41^ and the AMBER99-ILDN force field^42^ in combination with the TIP3P water model.^43^ The system was simulated in the NPT ensemble with periodic boundary conditions. The particle mesh Ewald method was used to account for long-range electrostatics, with a cutoff of 12 Å.^44^ A time step of 2 fs was used for all simulations after constraining the hydrogen stretching modes using the LINCS algorithm.^45^

The presented SWISH-X simulation comprises six parallel replicas, each with its defined scaling factor. All replicas include the OPES Multithermal component to span a selected temperature range. The scaling factors were uniformly distributed and ranged from 1.00 (non-scaled replica) to 1.5. The selected temperature range ranged from the chosen thermostat temperature, 300K, to 330 K. Benzene was included in the simulations as a cosolvent at a concentration of 1 M. The benzene molecules and the protein system were assigned to the same temperature coupling group. The simulation parameters were consistent with those used in the unbiased MD simulations. We included RMSD restraints to prevent potential unfolding of the protein in replicas with high scaling factors. The optimal upper wall value for the restraints were determined by analysing the RMSD fluctuations during the unbiased simulations. The SWISH-X simulation was run with GROMACS 2023 patched with PLUMED 2.9 the AMBER99-ILDN force field and TIP3P water model. A link to a detailed guide on how to set up a SWISH-X simulation is available at https://github.com/Gervasiolab/Gervasio-Protein-Dynamics/tree/master/swishX_bootcamp.

## Data availability

All code developed for the creation and analysis of CryptoBank, as well as the finetuning of the protein language model, is available at https://github.com/Gervasiolab/CryptoBank. The version of CryptoBank used for the analysis presented in this study is accessible via Zenodo at https://zenodo.org/records/15212595. An interactive web interface for exploring CryptoBank and predicting cryptic binding sites is available at https://farma-unites.unige.ch/en/gervasio-lab/pages/cryptobank.

## Acknowledgments

We are grateful to V. Oleinikovas, A. Kuzmanic, C. Estarellas, and R. Evans for useful discussions and assistance in developing a first prototype of the PDB pipeline. The authors acknowledge PRACE and the Swiss National Supercomputing Centre (CSCS) for large supercomputer time allocations on Piz Daint, project IDs: pr126, s1107, s1169, s1228. FLG acknowledges the Swiss National Science Foundation and Bridge for financial support (projects number: 200021_204795, CRSII5_216587 and 40B2-0_203628).

