## Supplementary Material for "CryptoBank: A Resource for the Identification and Prediction of Cryptic Sites in Proteins"

#### Classifier training

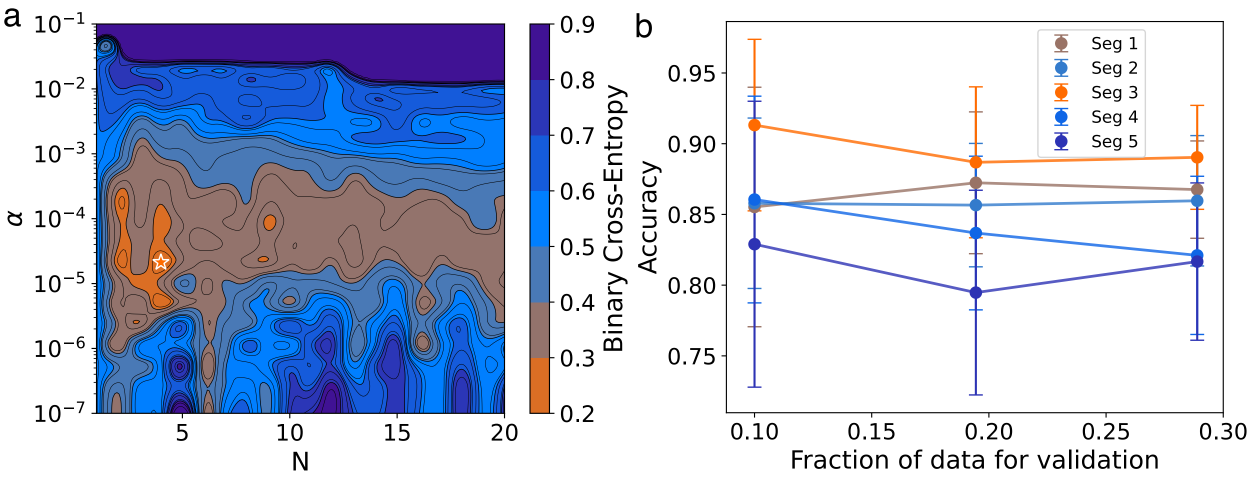


Figure S1: Hyperparameter scan over the number of shells, $N$ , regularization strength, $\alpha$. and ligand segments, $S$ (a) 2D landscape of binary cross-entropy loss for the classifier after training, evaluated on the validation set. The star marks the optimal combination of $N$ and $\alpha$ . (b) Validation accuracy as a function of the fraction of data left out for validation. The analysis compares classifiers trained with their respective optimal $N$ and $\alpha$ values, further examining performance across varying numbers $S$.

#### Outliers Identification

To investigate cases where individual ligands exhibit crypticity scores that deviate substantially from the average crypticity of the binding site, we computed a ligand-site deviation score for each ligand:

$$\Delta_i = \text{lig\_mean\_score}_i - \text{site\_mean\_score}_i$$

where $\text{lig\_mean\_score}_i$ is the average crypticity score of ligand $i$ within site $i$, and $\text{site\_mean\_score}_i$ is the average crypticity score of the site it binds. We limited this analysis to binding sites with more than one unique ligand, as such cases may highlight how ligands of different size and orientation can differentially expose cryptic and non-cryptic regions. To systematically detect extreme deviations, we normalized the deviation scores by calculating z-scores:

$$z_i = \frac{\Delta_i - \mu}{\sigma}$$

where $\mu$ and $\sigma$ are the mean and standard deviation of the $\Delta_i$ distribution, respectively. Ligands with $|z_i| > 3$ were considered outliers.

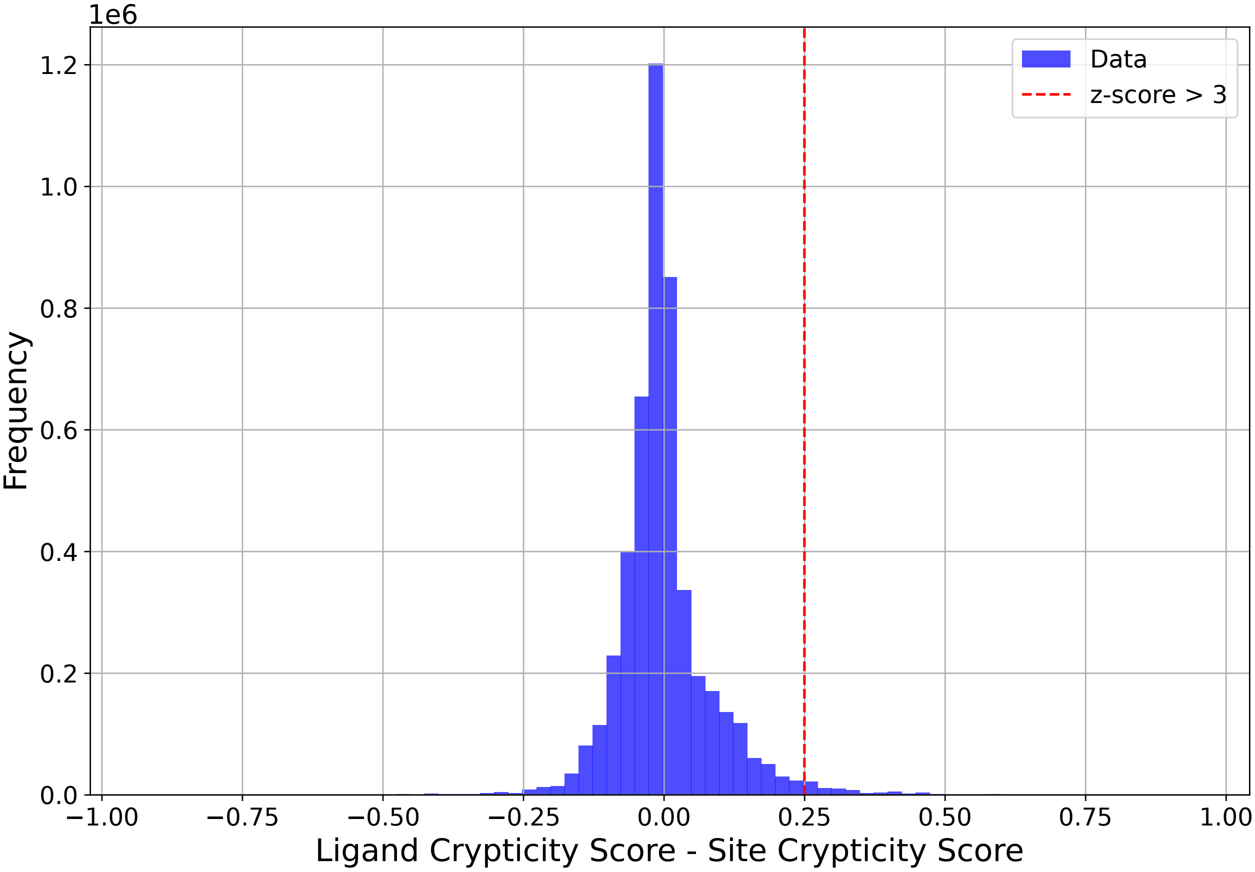


Figure S2: Histogram showing the distribution of the difference between the ligand crypticity score and the crypticity score of the site it binds to. The vertical dashed line indicates the threshold for right-tail outliers, defined as having a z-score greater than 3 ($z_i > 3$).

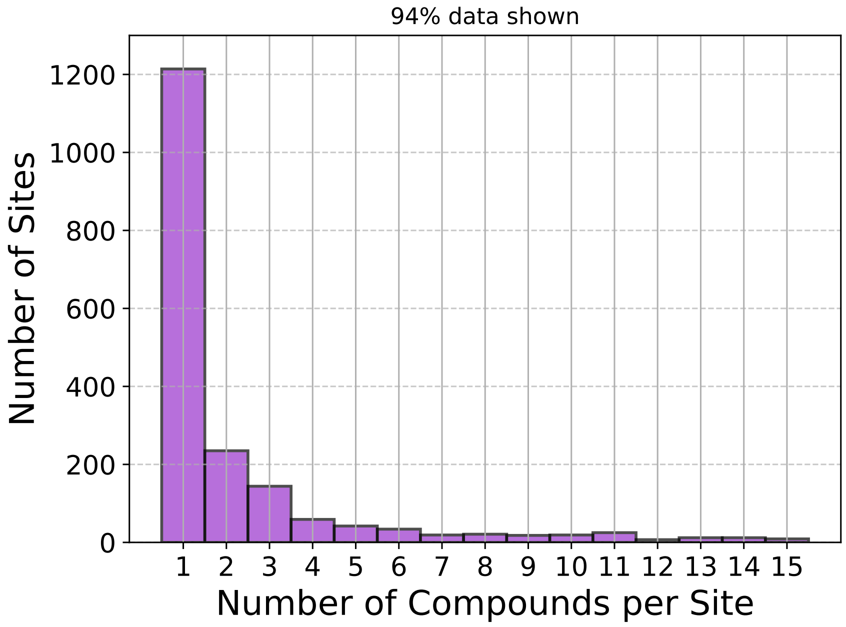


Figure S3: Histogram showing the distribution of the number of chemically distinct compounds binding to each cryptic site. The count is capped at 15 compounds per site, with sites binding up to 15 compounds cumulatively representing 94% of the total sites. Overall, 38.8% of cryptic sites bind more than one compound.

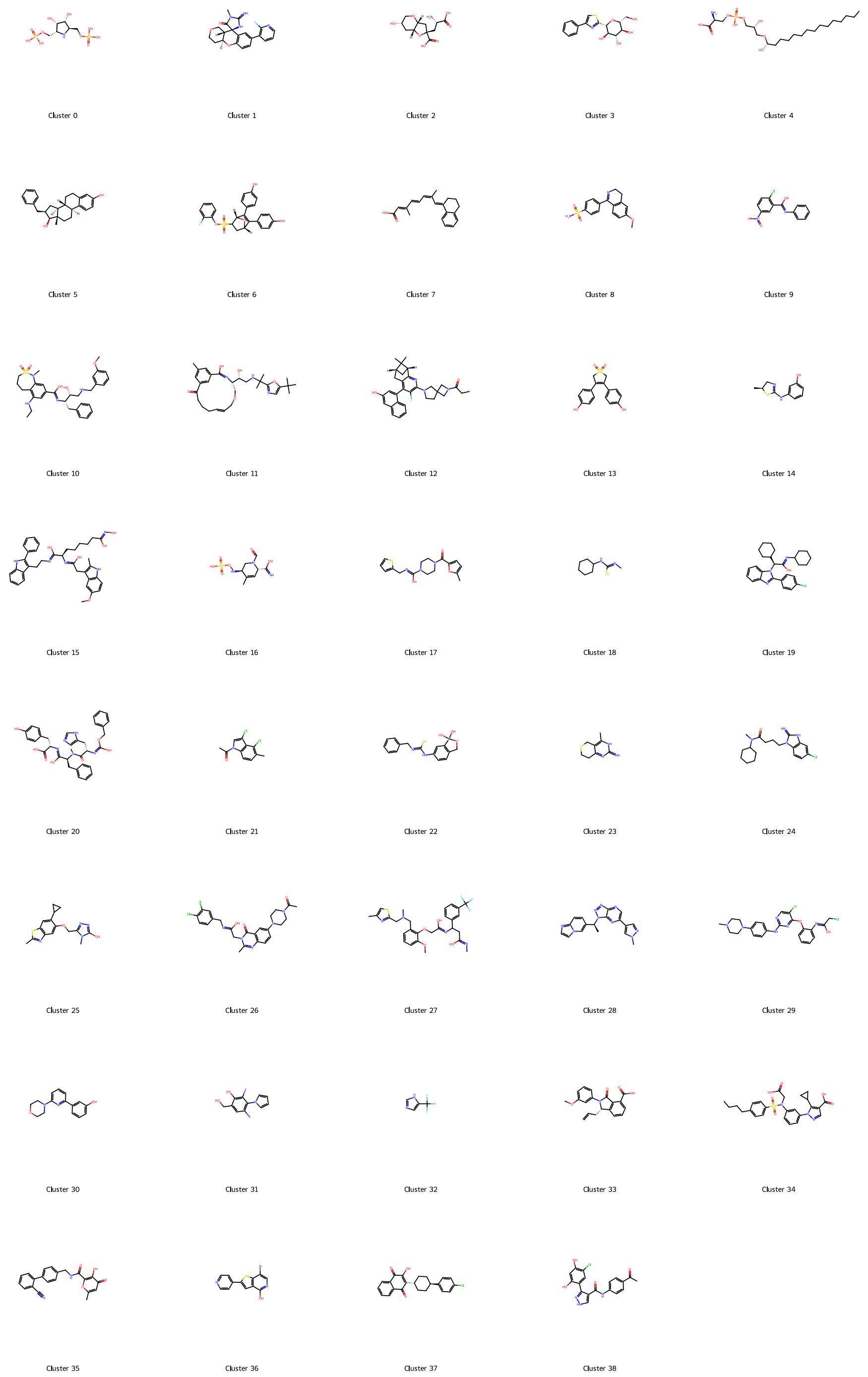


Figure S4: Chemical structures of the ligands selected as representatives for each cluster. The corresponding clusters are shown in Fig. S5a.

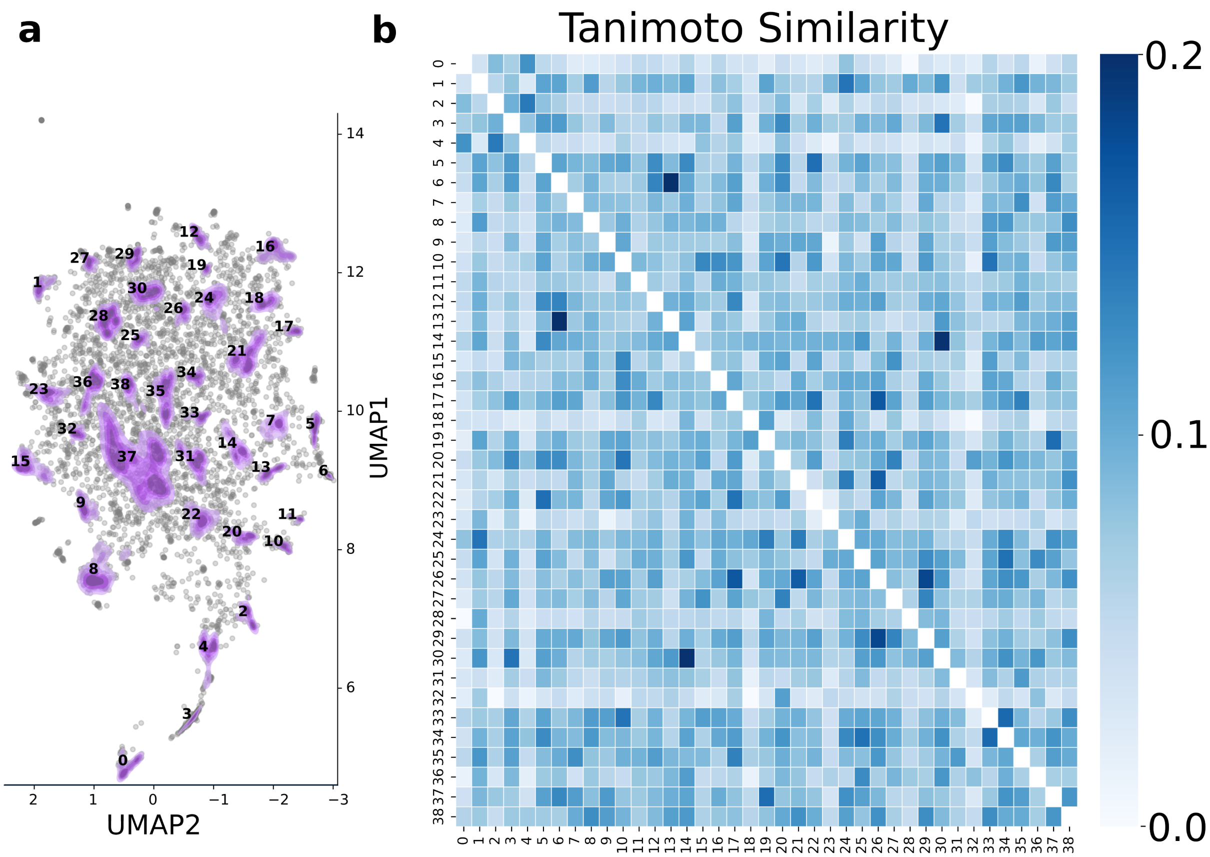


Figure S5: ***a)*** Chemical space of compounds found exclusively in cryptic binding sites, visualized using UMAP. Each point represents a compound, with clusters shown in purple and labelled numerically. Clustering was performed using HDBSCAN. Only compounds with a molecular weight below 1200 Da were included in the analysis. ***b)*** Heatmap showing the pairwise Tanimoto similarity between cluster centroids. The colour bar indicates the degree of chemical similarity.

### PLM fine-tuning for binding site prediction

In Figure S6 we collected the evaluation of the fine-tuned Prot-T5-XL-UniRef50 model performance across training, validation, and test set. While in the main text we show how PLM fine-tuning performs on exclusively cryptic sytems, this section highlights the extrapolation capabilities of a fine-tuned PLM focusing on binding site prediction, which effectively included more proteins and ligands. The ROC curve for the training set yields a high AUC of 0.96, because of the high data imbalance with a positive class ratio of 0.07 making it easier for the model to achieve high AUC. We select the best model as the one that is maximizing the cross-entropy loss on the validation set yielding an AUC of 0.89. Since the validation data share a similar imbalance with a positive class ratio of 0.06, the ROC AUC might be overestimate the extrapolation behavior to systems not seen during training. Benchmarking against the test set results in an AUC of 0.81 for the ROC curve (compare Figure S6(a)), indicating that unique sequences impose a significant challenge for the PLM, which is in agreement with the result of the fine-tuning experiment in the main text. To highlight the model's ability to achieve high precision we compute the model's precision-recall curves shown in Figure S6(b). According to this metric, the various datasets differ even more significantly in terms of their AUCs. The precision on the training data is high with a PR AUC of 0.80 corresponding to an increase in precision of an order of magnitude compared to a random model. Comparing to the precision of the selected model with respect to the validation set a significant decrease in PR AUC to 0.51 can be seen. Importantly the model retains significant precision indicating it is capable of extrapolation to new systems with a corresponding 9-fold increase in precision compared to random predictions. Benchmarking against the test set yielded an AUC 0.32 for the precision-recall curve. While the high accuracy obtained on the test set can be misleading for imbalanced data the PR AUC instead reveals the significant drop in performance. The retained precision corresponds to a 5-fold increase in precision compared to a random model.

The results of this second fine-tuning experiment indicate that signal about crypticity is encoded at the sequence level and that non-cryptic binding sites can be inferred from learning about cryptic ones. Furthermore, the model demonstrates a limited but notable ability to extrapolate signals to unique sequences, emphasizing its generalization capability beyond the training data.

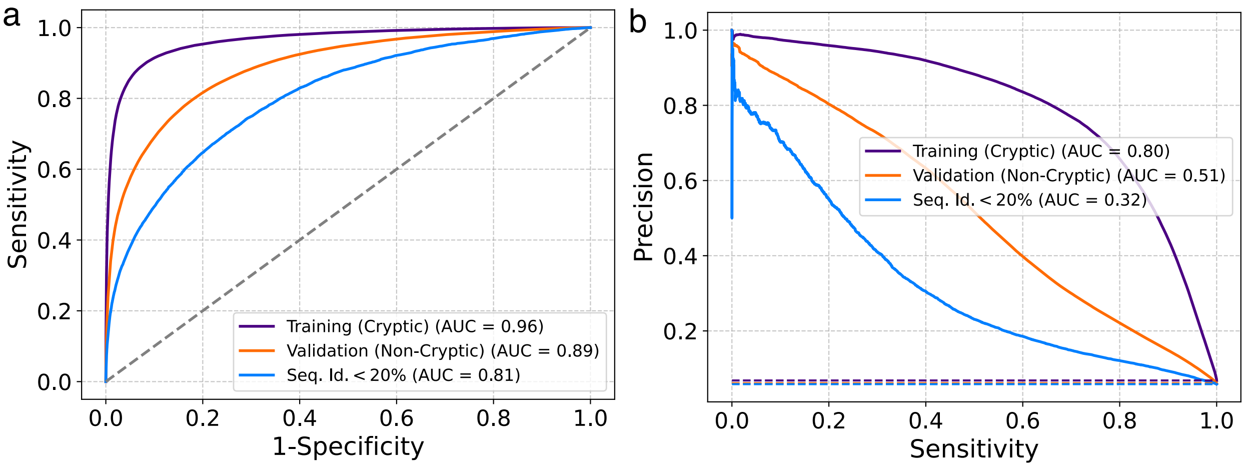


Figure S6: Performance of the 'Prot-T5-XL-UniRef50' model in predicting binding sites after fine-tuning evaluated on training, validation and test set. (a) ROC curves showing AUCs of 0.96, 0.89 and 0.81 respectively, while random prediction will lead to an AUC of 0.5. (b) Precision-recall curves show AUCs of 0.80, 0.51 and 0.32 respectively, demonstrating high precision for the imposed imbalanced binary classification task. Performance of random predictions corresponds to the fraction of the positive class and is maximally at 0.07 for all sets visualized as dashed lines with matching colors.
